# Which osteoarthritic gait features recover following Total Knee Replacement surgery?

**DOI:** 10.1101/398743

**Authors:** Paul Robert Biggs, Gemma Marie Whatling, Chris Wilson, Andrew John Metcalfe, Cathy Avril Holt

**Affiliations:** Cardiff School of Engineering, College of Physical Sciences, Cardiff University, Cardiff, UK; Arthritis Research UK Biomechanics and Bioengineering Centre, Cardiff University, Cardiff, UK; University Hospital of Wales, Cardiff, UK; Warwick Clinical Trials Unit, Warwick Medical School, University of Warwick, Coventry, UK

## Abstract

**Background:** Gait analysis can be used to measure variations in joint function in patients with knee osteoarthritis (OA), and is useful when observing longitudinal biomechanical changes following Total Knee Replacement (TKR) surgery. The Cardiff Classifier is an objective classification tool applied previously to examine the extent of biomechanical recovery following TKR. In this study, it is further developed to reveal the salient features that contribute to recovery towards healthy function.

**Methods:** Gait analysis was performed on 30 patients before and after TKR surgery, and 30 healthy controls. Median TKR follow-up time was 13 months. The combined application of principal component analysis (PCA) and the Cardiff Classifier defined 18 biomechanical features that discriminated OA from healthy gait. Statistical analysis tested whether these features were affected by TKR surgery and, if so, whether they recovered to values found for the controls.

**Results:** The Cardiff Classifier successfully discriminated between OA and healthy gait in all 60 cases. Of the 18 discriminatory features, only six (33%) were significantly affected by surgery, including features in all three planes of the ground reaction force (p<0.001), ankle dorsiflexion moment (p<0.001), hip adduction moment (p=0.003), and transverse hip angle (p=0.007). All but two (89%) of these features remained significantly different to those of the control group after surgery.

**Conclusions:** This approach was able to discriminate gait biomechanics associated with knee OA. The ground reaction force provided the strongest discriminatory features. Despite increased gait velocity and improvements in self-reported pain and function, which would normally be clinical indicators of recovery, the majority of features were not affected by TKR surgery. This TKR cohort retained pre-operative gait patterns; reduced sagittal hip and knee moments, decreased knee flexion, increased hip flexion, and reduced hip adduction. The changes that were associated with surgery were predominantly found at the ankle and hip, rather than at the knee.

## Introduction

Total Knee Replacement (TKR) surgery is a common procedure to treat late-stage knee osteoarthritis (OA), which aims to improve quality of life through the restoration of joint function and reduction of pain. Despite several studies reporting functional limitations following surgery, there appears to be a trend towards utilisation of TKR in younger patients with higher functional expectations [1–3]. The improvement of underlying joint biomechanics during gait is considered an important aspect of functional recovery following surgery [4] and is associated with post-operative activity levels [5].

There are numerous challenges to the adoption of three-dimensional gait analysis (3DGA) techniques within routine clinical assessment. Marker-based motion capture is typically considered infeasible when considering the resources required and volume of patients [6]. There are, however, several new measurement devices and assessment techniques currently being developed which may overcome many of these challenges[7]. With the potential of clinically feasible 3DGA techniques on the horizon, there is an increased importance in developing objective techniques to characterise the biomechanical outcome of surgery and communicate 3DGA findings to non-specialist audiences.

Longitudinal studies adopting 3DGA techniques have identified a number of abnormal biomechanical parameters which do not recover following TKR surgery [6,8,17,18,9–16]. Of these studies, many do not include biomechanical parameters of the hip and ankle [6,12–16], and/or exclude parameters within the frontal [6,14–17] or transverse [6,8,17,9–16] plane. There is compelling evidence that kinematic and kinetic changes are seen in knee OA subjects in all three planes of the hip, knee and ankle [19,20]. We believe that multivariate techniques that objectively characterise biomechanical outcomes in all three planes of the hip, knee and ankle will enhance our understanding of the biomechanical response to TKR as well as aid future development of multifactorial 3DGA outcome measures.

Instrumented lower-limb 3DGA results in a wealth of temporal waveforms. This vast dataset is then typically reduced into a considerably smaller set of discrete metrics (maximum, range, integral) calculated from selected waveforms. The measures must be defined *a priori* to avoid false-positive findings[21], and have been criticised for inherently disregarding the dynamic and highly collinear nature of biomechanical waveforms during motion [22]. Principal Component Analysis (PCA) is a multivariate technique that objectively defines features of variation from time-varying waveforms. The technique has the advantage of objectively described modes of variation across the entire waveform, often accounting for highly correlated features, such as peaks, loading rate, and range of motion within a single component [22–24].

In our unit, the application of PCA has been combined with a classification method based on a Dempster-Shafer Theory (DST) of evidence, termed the ‘Cardiff Classifier’. The principal application has been a summary gait measure, or index, which characterises the biomechanical changes associated with knee OA [25], and has used these features as an index to monitor recovery following TKR [10,26,27]. Metcalfe *et al.* expanded the initial application of this technique by including sagittal and transverse kinetics and kinematics of the ankle and hip[10]. Of the 17 biomechanical features found to be discriminatory between OA and non-pathological gait, seven were features of the hip or ankle, and only three were of parameters at the knee. Metcalfe *et al.* did not investigate which of these biomechanical features, if any, were significantly changed by TKR surgery and which remained significantly different from the non-pathological cohort.

The aim of this study is to identify which biomechanical features of OA significantly change following surgery, including features in the sagittal, frontal and transverse planes of the operative hip, knee and ankle. The first objective is to use the DST classification technique to identify the strongest discriminating features of severe OA vs non-pathological gait. The second objective is to test whether these features were significantly affected by TKR, and if so, whether they are normalised to that of a non-pathological cohort.

## Methods

### Study Participants

We performed a prospective, longitudinal study of a patient cohort with knee OA undergoing TKR surgery. The study was approved by the Research Ethics Committee for Wales and Cardiff and Vale University Health Board. Participants were excluded if they were unable to walk 10m without a walking aid, were unable to give informed consent, or had an unrelated musculoskeletal, neurological or visual condition that might affect the way they move. Participants were assessed pre-operatively and again at a target of 12 months post-operatively. At the time of analysis, 30 subjects had undergone post-operative assessment. Due to several practical issues, there was variability in the timing of follow-up visit – the median time was 13 months but this ranged between 8 and 26 months following surgery. An initial analysis confirmed there was no relationship between post-operative time-point and outcome assessed using the Oxford Knee Score (OKS).

We recruited 30 non-pathological (NP) volunteers into the study. The inclusion criteria matched that of TKR subjects, with the addition of no history of musculoskeletal conditions that required medical treatment, and no self-reported pain in the lower-limb or back.

On the day of biomechanical assessment, volunteers were asked to complete the OKS, which was scored ranging from 0 (worst outcome) to 48 (best outcome). OKS pain and function subscale scores were calculated following the method of Harris et al [28] the function subscale score is obtained by summing the scores for OKS questions 2, 3, 7, 11, and 12 and the pain subscale score by summing scores for the remaining seven questions. These are then represented as a percentage with 100% being the best outcome.

### Biomechanical Analysis

Human motion analysis was performed during level gait at the motion analysis laboratory at Cardiff School of Engineering. A lower-limb CAST marker set [29] was attached to subjects, while they walked barefoot at a self-selected pace along a 10m walkway. Marker trajectories were collected using eight Oqus (Qualisys, Sweden) cameras capturing at 60Hz, and Ground Reaction Forces (GRF) were calculated from two force platforms (Bertec, USA) capturing at 1080Hz. Hip, knee and ankle kinematics and kinetics were calculated within Visual 3D (C-Motion, USA).

### Data Reduction

PCA was performed on the waveforms of OA and NP subjects to define distinct biomechanical features of variation between and within the cohorts. We initially selected the first three Principal Components (PCs) of each input variable, resulting in 69 discrete variables per subject. Following the recommendations of Brandon *et al.* [24], we performed single component reconstruction alongside representative extremes of each PC to aid interpretation of the biomechanical feature reconstructed by each component. For ease of communication and interpretation, where the mean PC score for OA subjects was higher, the PC scores and eigenvectors for all groups were negated such that a low PC score always corresponded the feature associated with osteoarthritic function. This consistent sign convention has no further statistical effect within the analysis.

The Cardiff Classifier was then used to rank input variable importance. This ranking deviated from a previously reported method [10] to reduce the risk of over-fitting, the training data was split into two equal halves and the classifier was used to rank the input variables within both data sets. There were 18 biomechanical variables which were identified as being highly ranked in each group and were retained for further analysis.

### Data Classification

The 18 discrete biomechanical features, which discriminated between the 30 NP and 30 pre operative TKR subjects, were used to train the Cardiff Classifier on the characteristics of OA gait. This process defines the relationship between each of the input features, and a belief value of OA, NP and Uncertainty. These three belief values termed B(OA), B(NP) and Urespectively, are then used to classify between OA and NP gait biomechanics [26]. If, for example, B(OA) is greater than B(NP), and the subject belongs to the OA group, the classification technique is deemed to have successfully classified this subject. We addressed the robustness of this classification using the leave-one-out (LOO) cross-validation algorithm.

We then applied the same process to the lower-limb biomechanics collected at the follow-up visit, using the previously defined PCs to calculate scores for the 30 subjects following surgery. The same 18 biomechanical features were also inputted into the trained classifier to calculate the three belief values B(OA), B(NP) and U at the follow-up visit.

### Statistical Analysis

Paired samples tests were carried out within MATLAB Statistics and Machine Learning Toolbox (MathWorks, USA) to test for significant changes following surgery. Where parametric assumptions were not met, a Wilcoxon signed rank test was used. A t-test was used to identify differences between the post-operative TKR and the NP group. Where parametric assumptions were not met, the Mann-Whitney test was used. A Bonferonni correction was used to adjust for multiple comparisons. All statistical inferences were calculated using the MATLAB Statistics and Machine Learning Toolbox (MathWorks, USA).

## Results

Participant characteristics are summarised in Table 1. The TKR cohort was significantly older and had a higher Body Mass Index (BMI) than the NP control participants. The OKS was significantly improved following surgery by a mean(SD) change of 14.7(8.8) points, however, it remained significantly worse than that of NP subjects. There were significant improvements in both OKS pain and function subscales, with greater improvements seen in the OKS pain score. Gait velocity increased significantly following surgery but remained significantly lower than NP controls following TKR.

**Table 1:** Differences in clinical characteristics and principle component scores of kinematic and kinetic waveforms between the pre-surgery, post-surgery and between the non-pathological and post-surgical group.

| Parameters | Pre- TKR<br>Mean (SD) | Post- TKR<br>Mean (SD) | NP<br>Mean (SD) | P-value |  |
| --- | --- | --- | --- | --- | --- |
|  |  |  |  | pre-post | NP-post |
| Sex (F/M) | 15F, 15M |  | 18F, 12M |  |  |
| Age (y) | 69.7 (8.6) | 70.7 (8.3) | 39.8(17.6) |  | p<0.001* |
| BMI (kg/m<sup>2</sup>) | 32.5 (6.5) | 33.3 (6.9) | 24.3(3.7) | 0.2737 | p<0.001* |
| Oxford Knee Score | 19.5 (9.0) | 34.6 (9.9) | 48.0 (0.0) | p<0.001* | p<0.001* |
| OKS Pain<sup>a</sup> (%) | 27.5 (20.0) | 72.3 (26.2) | 100 (0.0) | p<0.001* | p<0.001* |
| OKS Function<sup>a</sup> (%) | 45.3 (18.7) | 65.3 (19.1) | 100 (0.0) | 0.002* | p<0.001* |
| Gait velocity (m/s) | 0.73 (0.21) | 0.87 (0.21) | 1.24 (0.12) | p<0.001* | p<0.001* |

\*statistically significant, p<0.01
NP = Non-pathological, OKS = Oxford Knee score, SD = Standard deviation, TKR = Total Knee Replacement.

The Cardiff Classifier was able to correctly classify between NP and OA gait biomechanics in all 60 cases, assessed using the LOO cross-validation technique. The three belief values are shown in a simplex plot within Fig 1. One pre-TKR subject was close to the decision boundary and had the second highest pre-operative OKS of 34/48.

**Figure 1:**
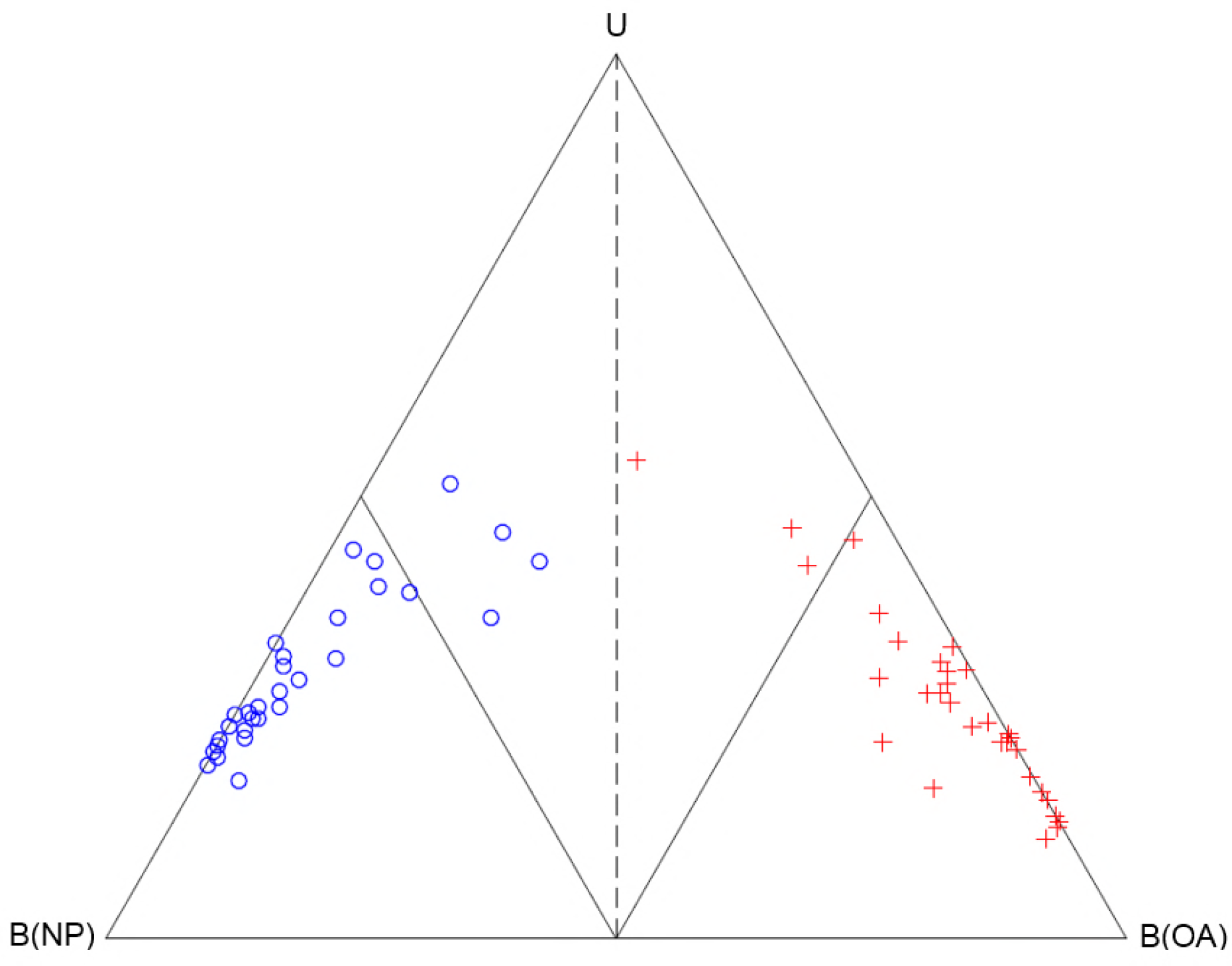
Simplex plot of the classification of the 30 NP (blue circle) and 30 pre-TKR (red cross) subjects which were used to train the Cardiff Classifier on the biomechanical features of severe osteoarthritic gait. The three vertices represent the points where belief of non-pathological function B(NP), belief of osteoarthritic function B(OA) and uncertainty, U is equal to 1 (or 100%). The decision boundary where B(OA)=B(NP) is shown as a dashed line. The boundaries where B(OA)=0.5 and B(NP)=0.5 are shown as interior solid lines.

There were 18 PCs retained for analysis; their accuracy in discriminating OA gait is displayed within Table 2. Also shown is the interpretation of the biomechanical feature, which is represented by each PC. The single-component reconstructions for the NP and OA subjects are displayed in supporting figures S1-S4. The greatest accuracy (100%) was achieved using PC1 of the vertical GRF.

**Table 2.**
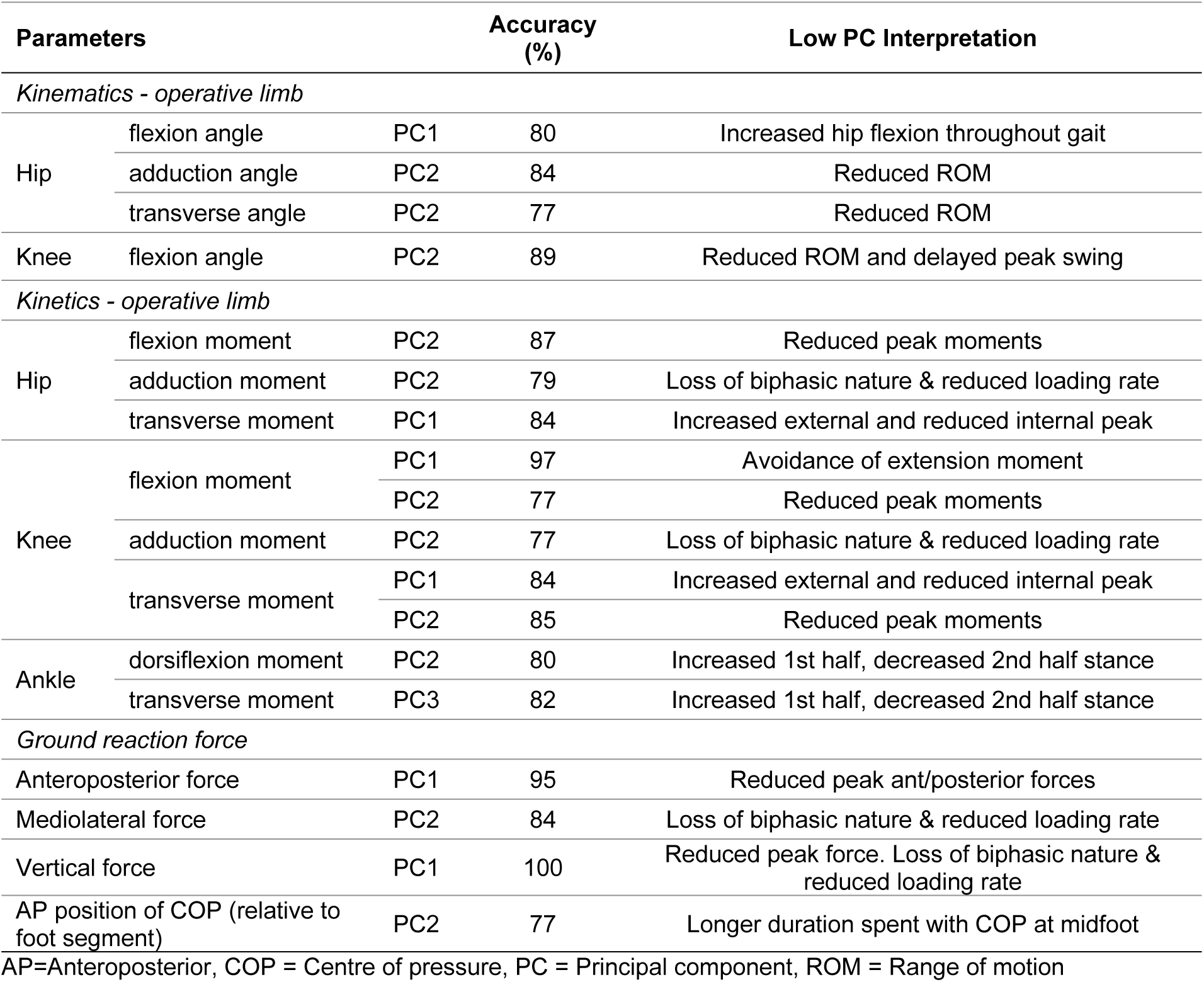
Classification accuracy of each input variable within the classifier, and the interpretation of the biomechanical feature represented by a low PC score.

The change in belief values following the TKR, relative to the pre-operative assessment, is shown in Figure 2. Only three subjects returned towards the healthy side of the classifier, 16 subjects remained in the “dominant” OA region where B(OA)>0.5, and one subject saw a decline in function from the non-dominant to the dominant region.

**Figure 2:**
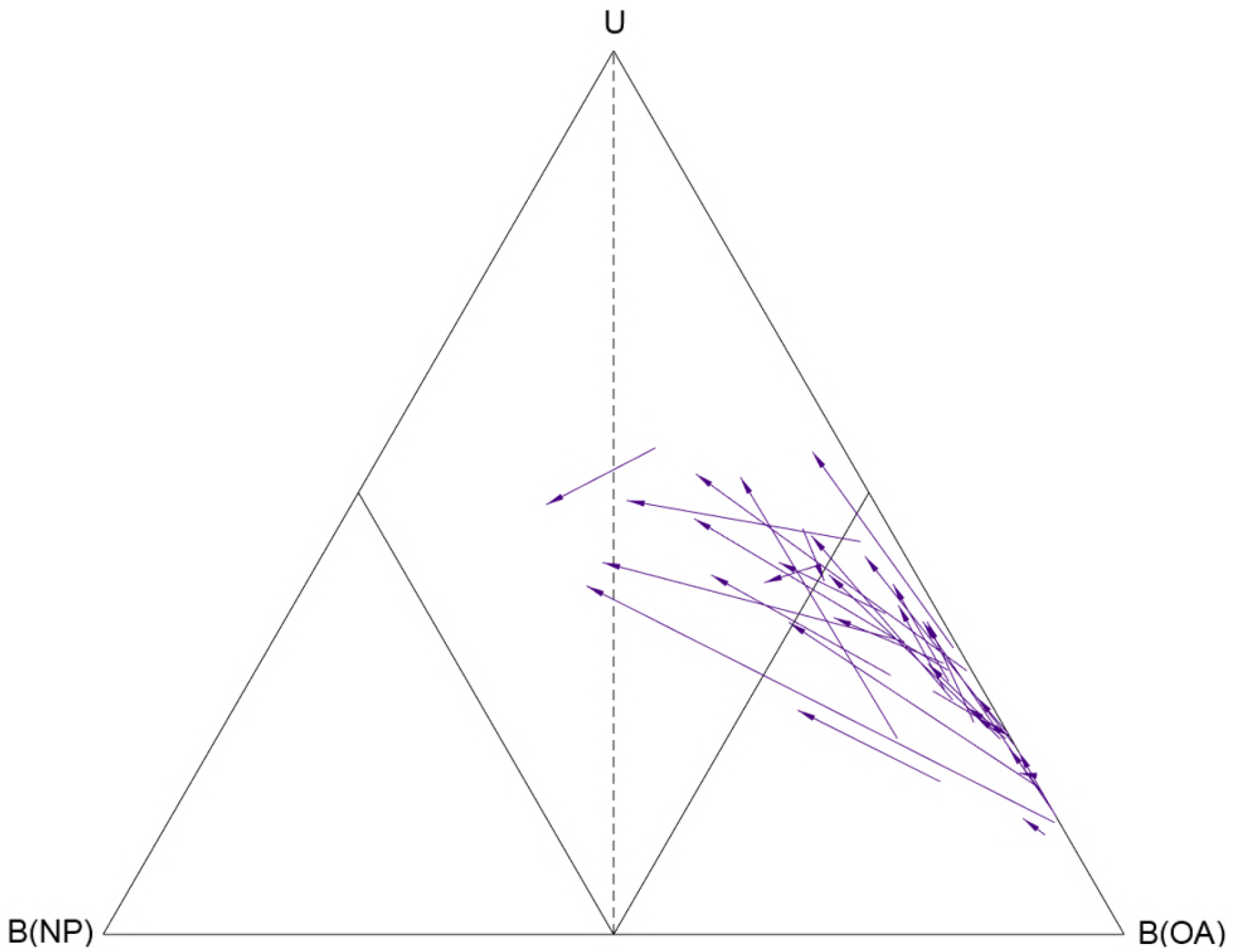
Simplex plot of the change in classification of the 30 TKR subjects between pre-and post-operative visits. The three vertices represent the points where belief of non-pathological function B(NP), belief of osteoarthritic function B(OA) and uncertainty, U is equal to 1 (or 100%). The decision boundary where B(OA)=B(NP) is shown as a dashed line. The boundaries where B(OA)=0.5 and B(NP)=0.5 are shown as interior solid lines. The purple arrows represent the change in the body of evidence for each subject from the pre-operative visit (arrow tail), to the post-operative visit (arrow head).

The changes in the individual biomechanical features (PCs) are within Table 3. Significant improvements following surgery were observed in only 6 of 18 features, and 15 features remained significantly different to the NP cohort post-operatively. Improvements were measured in all three planes of the GRF, alongside the transverse hip angle, hip adduction moment, and the ankle dorsiflexion moment. None of the six biomechanical features of the knee selected for analysis saw significant improvements following surgery. Moderate improvements were seen in PC2 of the knee flexion angle and flexion moment but these were not significant following Bonferroni correction.

**Table 3.**
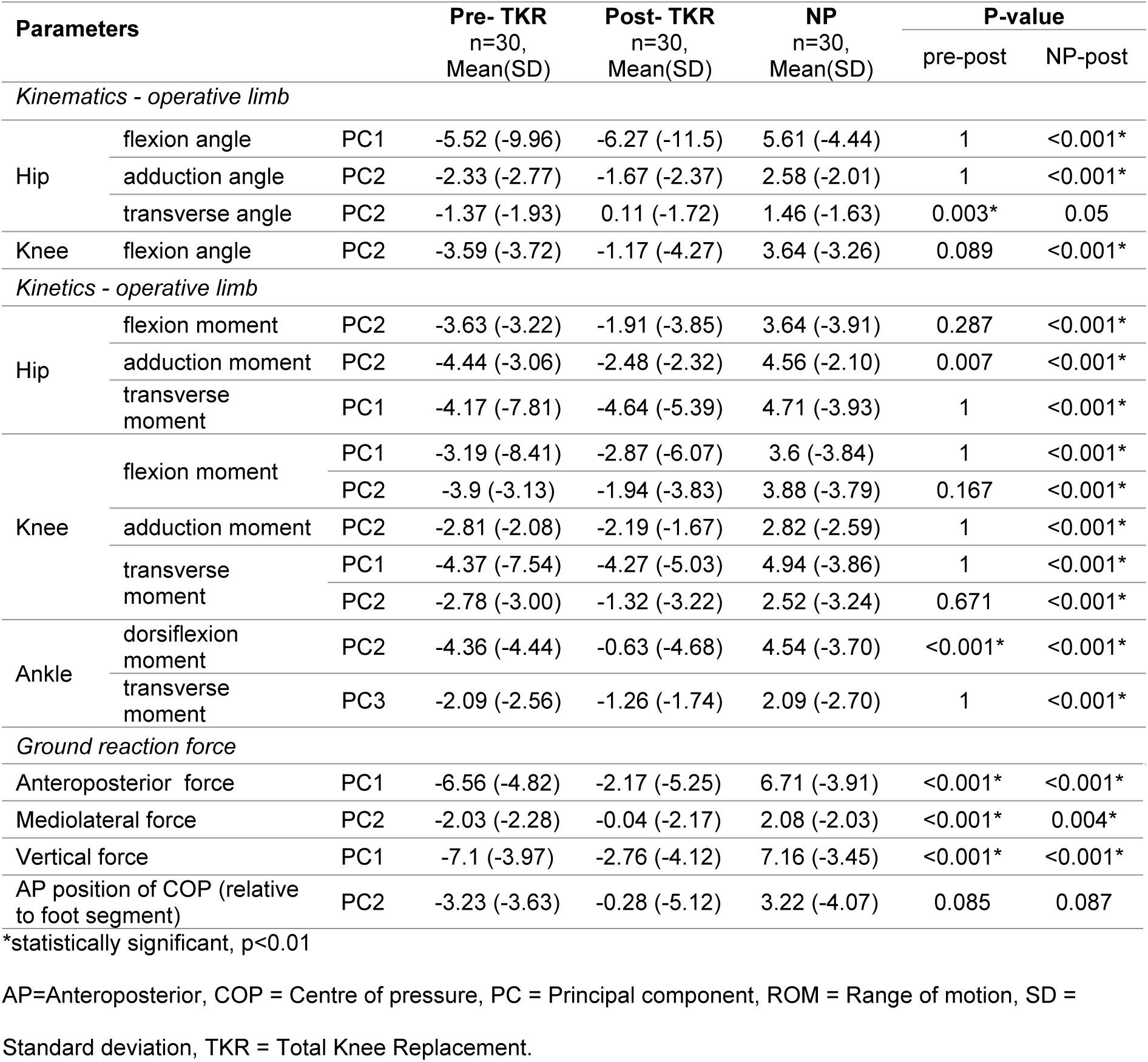
Differences in principle component scores of kinematic and kinetic waveforms between the pre-surgery, post-surgery and between the non-pathological and post-surgical group.

## Discussion

The biomechanical function of TKR subjects within this study did not return to that of the NP cohort. Of the 18 biomechanical features which all have >75% accuracy in discriminating OA gait within the NP cohort, significant improvements were only observed in six features, and none of the six retained features at the knee saw significant improvements. Considering the OKS thresholds proposed by Hamilton[30], 20 of 29 patients (1 OKS missing) achieved a successful outcome. Out of these nine subjects with a poor subjective outcome, six had remained in the “dominant” region of the simplex plot, where B(OA)>0.5.

Key to the interpretation of the biomechanical findings is that the gait velocity of TKR patients did not return to that of the NP cohort. Significant associations between gait velocity and numerous biomechanical parameters have been highlighted within both NP and pathological gait [31]. Several studies correct for this by considering gait velocity as a covariate within the statistical analysis, however, this violates the primary assumption that the co-variate isn’t related to the main effect [31]. Instead, we chose not to control speed and present un-altered data and to accept that the causal relationship between biomechanical changes and gait velocity cannot be determined within this study. This is typical of numerous similar studies [9– 13,15].

The control group NP cohort were significantly younger, with a lower BMI than the patient cohort. As biomechanics are affected by both age and BMI, the normalisation of parameters to that of the NP control may not be a realistic goal following surgery. Our decision not to age or BMI-match reflects the desire to exclude those who either have or are at high risk of developing musculoskeletal conditions which might affect hip, knee or ankle biomechanics. Both ageing and obesity are risk factors of OA, alongside a number of other comorbidities that affect locomotion[32]. Furthermore, a recent meta-analysis indicates that prevalence of knee OA features in asymptomatic adults increases linearly with age with approximately 75% of adults aged >70 having a cartilage lesion [33].

Features extracted from the GRF were a strong discriminator of severe OA function and showed significant improvement following TKR surgery. This is interesting considering this data is by far the least challenging and most clinically feasible to extract and process. Previous studies have highlighted the ability to discriminate pathological function from GRFs [34–36] and have suggested its use as an outcome measure following intervention[34]. Parameters of the vertical GRF commonly defined in other studies, such as loading rate and peaks during weight acceptance and push off, alongside the ratio of the peaks to the trough at midstance, are all represented in a single feature. This indicates high collinearity between these features.

Similarly to the findings of other studies, the second PC of the knee flexion angle was a better discriminator of OA than PC1, despite accounting for only 24% of the total variance [17,23]. The variance reconstructed by this PC is also very similar in our study: reduced Range of Motion (ROM) during stance phase and a reduced and delayed peak flexion during swing phase. Changes following surgery did not reach statistical significance and remained significantly different from the NP cohort following surgery. Ro *et al.* observed a much larger change in PC1 of the knee flexion angle following TKR, which represented a magnitude offset throughout the waveform. Although not retained during feature selection, we tested PC1 within our dataset and found no significant difference even before post-hoc correction. Restoration of sagittal knee kinematics during gait is an important functional goal following surgery which has not been met within this cohort.

The first PC of the hip flexion angle, representing increased flexion throughout the gait cycle, was also a highly-ranked discriminator of OA gait. Both decreased hip flexion and increased anterior pelvic tilt has been reported in elderly and OA gait [37–39]. This feature was not affected by TKR and therefore remains significantly different following surgery. It is possible that increased hip flexion could have been a strategy to increase ground clearance in the presence of insufficient knee ROM. Ouellet and Moffet reported increased hip flexion two months after TKR and suggested it may form a strategy to compensate for weak quadriceps [11]. It is, however, also possible that increased hip flexion in our cohort was a consequence of increased pelvic tilt. We recommend future work reports on both the angle of the pelvis and angle of the thigh segment in relation to the laboratory floor to elucidate the underlying mechanism of this gait alteration.

Frontal and transverse hip kinematics are also abnormal pre-operatively, with a significant improvement in hip internal/external angle PC1, and no improvement in hip adduction PC2 following surgery. Both PCs reconstruct changes in ROM through the gait cycle. During healthy gait, the pelvis typically drops a small amount towards the leg in swing phase. This movement results in increased pelvic obliquity and hip adduction of the leg in stance, and is exaggerated in the presence of hip pathology [40]. The second PC of the hip adduction angle, however, appears to show a reduction of this mechanism. Interestingly, Liebensteiner *et al.* previously identified a ‘paradoxical’ positive relationship between pelvic obliquity during stance and knee function [41]. One possible explanation for these findings is that knee OA and TKR with inferior knee function adopt a strategy known as hip hiking [42], perhaps as a compensatory mechanism to increase ground clearance in the presence of insufficient knee or hip flexion.

Frontal plane kinetics were consistent with numerous other studies which highlight the reduction in the “biphasic” nature of frontal plane joint moments due to OA [23], which remain following TKR. The second PC of the hip and knee adduction moments reconstructed very similar features, however, improvements were only observed at the hip following TKR. A ‘flat’ knee adduction moment both before and after surgery, where two peaks are not clearly identifiable, can also be observed in several other studies [9].

Sagittal and transverse plane kinetics were consistent with changes associated with reduced gait velocity. Retained PCs in the sagittal and transverse planes of the hip and the sagittal plane of the knee represent reduced joint moments at loading response and push off, consistent with the observed reduction in the Anteroposterior (AP) GRF. Interestingly, despite a significant increase in gait velocity and AP force following TKR, sagittal features of the hip and knee were not significantly improved. A possible explanation is that an increased gait velocity was more strongly related to changes in the ankle, as opposed to the hip and knee. This certainly seems consistent with the significant improvement in PC2 of the ankle plantarflexion moment observed following surgery.

The retained PC of the plantarflexion moment is challenging to interpret and requires the consideration of PC1, which was not retained for further analysis. PC1 represented 46% of the variance and reconstructs changes in the magnitude of the waveform from loading response to push off. In comparison, the second PC reconstructs a similar reduction towards push off, however, this is related to an increased moment during the first half of stance. While accounting for less variance (36%), PC2 was more characteristic of changes relating to OA. These findings are corroborated by the differences detected in the Centre of Pressure (COP) of the GRF relative to the foot during the stance phase. While post-operative changes in PC2 of the AP position of the COP did not reach significance, this feature was no longer significantly different from that of NP subjects. The PC shows that the COP progresses faster towards the mid-foot in early stance, and faster towards the forefoot in late stance. Relating to the “three rockers” described by Perry [43], OA subjects progressed faster toward the ankle rocker, where the foot is typically flat on the ground.

One of the limitations of this study is that the limited sample size (n=30), in comparison to the number of statistical inferences made, could increase the chance of erroneous results. A Bonferroni correction was therefore applied, which is generally considered a conservative approach. The small inclusion criteria for the patient cohort of this study increases the generalisability of these findings, however, the resultant cohort is heterogeneous with numerous comorbidities. It is, therefore, possible that differences in biomechanical outcomes between different clinical phenotypes of OA are masked when treating these phenotypes as a single group. A further limitation was the large range in time-points of the post-operative visits (range 8-26 months). While such a broad range in time-points is common in studies assessing post-operative TKR biomechanics [4], there is some evidence of functional [44] and biomechanical changes across this timeframe [45].

We consider the inclusion of all three planes of the hip, knee, and ankle within the description of the biomechanical change to be a strength of this study. The presented method of reducing the included 23 waveforms, all normalised to 101 data points, to a subset of 18 biomechanical features, is both objective and generic.

## Conclusion

This study found that most biomechanical features of OA were not significantly normalised to that of the NP control cohort following TKR surgery. Furthermore, despite improvements in reported pain and function, only 6 of 18 identified discriminatory features were affected by surgery: two features of the hip, one of the ankle, three of the GRF, and none of the knee. We observed no effect of TKR on sagittal knee and hip kinetics despite an increase in velocity. TKR patients maintained reduced knee flexion, which may relate to increased hip flexion and frontal plane hip hiking during gait. The identified discriminatory features may be good targets for assessing outcome in future studies. Notably, the GRF is both the easiest to measure and has the strongest discriminatory effect.

